# CD38 mediates nicotinamide mononucleotide (NMN) base exchange to yield nicotinic acid mononucleotide (NaMN)

**DOI:** 10.1101/2024.08.08.607247

**Authors:** Romanthi Madawala, Jasmine L. Banks, Sarah E. Hancock, Lake-Ee Quek, Nigel Turner, Lindsay E. Wu

## Abstract

Nicotinamide mononucleotide (NMN) is a widely investigated metabolic precursor to the prominent redox cofactor nicotinamide adenine dinucleotide (NAD^+^), where it is assumed that delivery of this compound results in its direct incorporation into NAD^+^ via the canonical salvage / recycling pathway. Surprisingly, treatment with this salvage pathway intermediate leads to increases in nicotinic acid mononucleotide (NaMN) and nicotinic acid adenine dinucleotide (NaAD), two members of the Preiss-Handler / *de novo* pathways. In mammals, these pathways are not known to intersect prior to the production of NAD^+^. Here, we show that the cell surface enzyme CD38 can mediate a base exchange reaction on NMN, whereby the nicotinamide ring is exchanged with a free nicotinic acid to yield the Preiss-Handler / *de novo* pathway intermediate NaMN, with *in vivo* small molecule inhibition of CD38 abolishing the NMN-induced increase in NaMN and NaAD. Together, these data demonstrate a new mechanism by which the salvage pathway and Preiss-Handler / *de novo* pathways can exchange intermediates in mammalian NAD^+^ biosynthesis.

## Introduction

Nicotinamide adenine dinucleotide (NAD^+^) is a prominent redox cofactor that declines during ageing and physiological challenges such as DNA repair and inflammation. Over the past decade, there has been strong interest in restoring NAD^+^ levels as a therapeutic approach to overcoming age-related diseases including frailty, infertility, and impaired muscle function (Aflatounian, Paris et al., 2022, Bertoldo, Listijono et al., 2020, Das, Huang et al., 2018, Rajman, Chwalek et al., 2018). These approaches have included the use of the metabolic precursors nicotinamide riboside (NR) (Mouchiroud, Houtkooper et al., 2013, Wu & Sinclair, 2016, Zhang, Ryu et al., 2016) and nicotinamide mononucleotide (NMN) (Mills, Yoshida et al., 2016), both of which are amidated precursors that act as substrates for NAD^+^ biosynthesis via the salvage or recycling pathways – in the case of NR, first entering pathway following its phosphorylation via the NRK pathway (Belenky, Christensen et al., 2009, Bieganowski & Brenner, 2004, Tempel, Rabeh et al., 2007). The other well-characterised pathways to NAD^+^ biosynthesis in mammals are the Preiss-Handler pathway, which utilises nicotinic acid as a substrate for the production of nicotinic acid mononucleotide (NaMN) (Preiss & Handler, 1958a, Preiss & Handler, 1958b), and the *de novo* pathway, which converts the amino acid tryptophan into NaMN, both of which are converted into nicotinic acid adenine dinucleotide (NaAD) prior to the final step of its conversion to NAD^+^.

One core chemical difference between these pathways is that the salvage pathway utilises amidated intermediates, whereby the nicotinamide ring contains an amine group as a sidechain from the pyridine ring (Fig. 1). In contrast, the Preiss-Handler / *de novo* pathways utilise intermediates containing nicotinic acid, which contains a carboxylic acid sidechain, and could be considered as deamidated counterparts to amidated precursors in the salvage pathway. In mammals, these pathways do not intersect, and there are no mammalian enzymes known to mediate the deamidation of salvage pathway intermediates into their deamidated counterparts (Fig. 1A). Despite this, administration with the amidated intermediates NR and NMN leads to striking increases in the deamidated metabolites NaMN and NaAD (Igarashi, Nakagawa-Nagahama et al., 2022, Kim, Chalmers et al., 2023, Trammell, Schmidt et al., 2016). The question of how treatment with the amidated intermediate NMN leads to an increase in levels of the deamidated intermediates NaMN and NaAD (Igarashi et al., 2022, Kim et al., 2023) is the core goal of this investigation, as it could point to overlap in these pathways, and provide an update to these textbook pathways of NAD^+^ biosynthesis.

**Figure 1.**
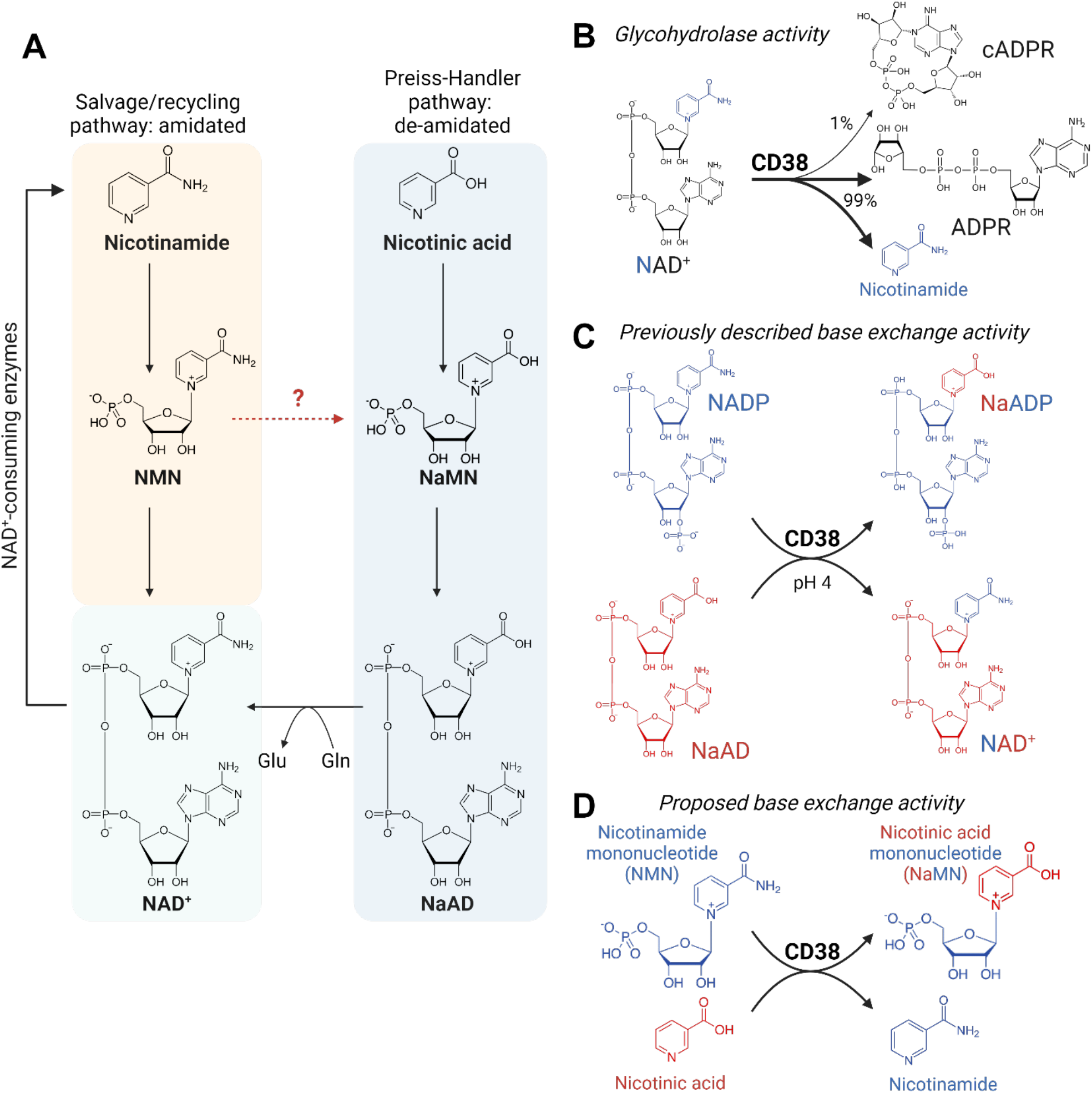
Mammalian NAD^+^ biosynthesis, and proposed role for CD38. A) NAD^+^ can be synthesised via the salvage pathway, which recycles the amidated precursor nicotinamide via an amidated intermediate, nicotinamide mononucleotide (NMN). In the Preiss-Handler pathway, nicotinic acid is incorporated into NAD^+^ via the deamidated intermediates nicotinic acid mononucleotide (NaMN) and nicotinic acid adenine dinucleotide (NaAD). Treatment with the amidated precursor NMN can increase levels of the deamidated intermediates NaMN and NaAD. B) The enzyme CD38 is well-studied for its NAD^+^ glycohydrolase activity, cleaving NAD^+^ into free nicotinamide and ADP-ribose (ADPR). C) CD38 also has base exchange activity towards NADP^+^, yielding the Ca^2+^ signalling intermediate NaADP^+^. D) This investigation identified NMN and nicotinic acid as new base exchange substrates for CD38, yielding NaMN and explaining the crossover between the amidated salvage/recycling pathways and the deamidated Preiss-Handler / de novo pathways of NAD^+^ biosynthesis.

One mechanism for the link between these amidated and deamidated pathways of NAD^+^ biosynthesis is a role for the gut microbiome in mediating the deamidation of these precursors when they are delivered orally (Chellappa, McReynolds et al., 2022, Kim et al., 2023, Shats, Williams et al., 2020), due to the expression of bacterial deamidase enzymes that are lacking in mammals, such as the enzyme PncC, which deamidates NMN into NaMN (Galeazzi, Bocci et al., 2011). Recently, we used a stable isotope tracing strategy to show that orally delivered NMN could be incorporated into NAD^+^ via the deamidated *de novo* pathway, and that incorporation via this pathway was reduced following ablation of the gut microbiome through antibiotics treatment (Kim et al., 2023). While these data showed a reduction in NMN incorporation via the NAD synthase (NADS) dependent deamidated pathway during antibiotics treatment, this did not lead to a complete ablation of incorporation via this route. Further, early results from the field suggested the existence of mammalian enzymes that could convert NMN into NaMN (Petrack, Greengard et al., 1965, Sarma, Rajalakshmi et al., 1961). Another complication is that previous results from ourselves and others using stable isotope labelled precursors have identified only a small fraction of double labelled NR or NMN being incorporated into NAD^+^ intact (Chellappa et al., 2022, Kim et al., 2023, Liu, Su et al., 2018), with the majority of NAD^+^ labelling only containing a single label from the ribose or nicotinamide groups, suggesting that these precursors could undergo gastrointestinal metabolism into free nicotinamide prior to their delivery and incorporation into NAD^+^ in other tissues.

One additional explanation for these observations could be that in addition to microbial deamidation (Chellappa et al., 2022, Kim et al., 2023, Shats et al., 2020), the amidated and deamidated pathways of NAD^+^ biosynthesis could intersect via base exchange, rather than deamidase activity, whereby the entire nicotinamide ring would be exchanged for a nicotinic acid. This was recently described for the membrane-bound enzyme BST1/CD173, which could convert NR into nicotinic acid riboside (NaR) via exchange of the entire nicotinamide ring for a nicotinic acid (Yaku, Palikhe et al., 2021). This has also been described for the NAD^+^ glycohydrolase SARM1, which can mediate base exchange on NAD^+^ with nicotinic acid and other similar compounds (Angeletti, Amici et al., 2022). The cell surface enzyme CD38 is well-studied for its role as an NAD^+^ glycohydrolase (Fig. 1B) in inflammation, infertility, senescence and ageing (Camacho-Pereira, Tarrago et al., 2016, Chini, Peclat et al., 2020, Peclat, Agorrody et al., 2024, Perrone, Ashok Kumaar et al., 2023, Tarrago, Chini et al., 2018), and there is strong interest in the use of small molecule CD38 inhibitors as a strategy to preserve NAD^+^ levels (Perrone et al., 2023, Tarrago et al., 2018). In addition to its glycohydrolase activity, CD38 also has base exchange activity (Fig. 1C), whereby it uses NaAD as a nicotinic acid donor for base exchange with nicotinamide adenine dinucleotide phosphate (NADP^+^) to yield the Ca^2+^ mobilising signalling intermediate, nicotinic acid adenine dinucleotide phosphate (NaADP) in endolysosomes (Aarhus, Graeff et al., 1995, Chini, Beers et al., 1995, Fang, Li et al., 2018, Li & Wu, 2021, Nam, Park et al., 2020). CD38 also has prominent aeinity towards NMN, acting as a glycohydrolase for its breakdown into free nicotinamide (Sauve, Munshi et al., 1998), and is capable of carrying out base exchange reactions on NMN for drug-like heterocyclic substrates (Preugschat, Tomberlin et al., 2008).

Given both its affinity for NMN (Sauve et al., 1998), and its base exchange activity towards other NAD^+^ metabolites (Aarhus et al., 1995, Nam et al., 2020, Preugschat et al., 2008), we hypothesised CD38 could mediate base exchange activity towards NMN (Fig. 1D), helping to resolve the question of how the oral administration of the amidated precursor NMN can lead to sharp increases in the deamidated precursors NaMN and NaAD (Igarashi et al., 2022, Kim et al., 2023). Here, we demonstrate that small molecule inhibition of CD38 abolishes the spike in NaMN and NaAD caused by NMN treatment, and further show that this is due to direct base exchange activity between NMN and free nicotinic acid, with similar activity towards a series of other nicotinic acid analogues. These findings demonstrate an important link between the amidated salvage pathway and the deamidated Preiss-Handler / *de novo* pathways for NAD^+^ synthesis in mammals.

## Results

In the current investigation, we sought to identify additional mechanisms through which exogenous treatment with the amidated NAD^+^ precursor NMN could elevate NaMN and NaAD levels (Igarashi et al., 2022, Kim et al., 2023), which also occurs during treatment with NR (Trammell et al., 2016). Our previous investigations (Kim et al., 2023) focused on the gain or loss of isotope labels on the amide group of NAD^+^ in animals treated with or without antibiotics, where we had sought to identify whether there was deamidation of this group by non-mammalian enzymes expressed by the gut microbiome. Here, we sought to investigate whether the formation of NaMN and NaAD following NMN treatment could also occur independently of direct deamidation. Instead, we deduced that if NMN could undergo endogenous base exchange with a free nicotinic acid, this mechanism could also explain the formation of NaMN and NaAD.

We had hypothesised that if base exchange on NMN could occur, that one candidate would be the membrane-bound enzyme CD38, which has been the subject of intense study in the field of NAD^+^ biology due to its role as an NAD^+^ glycohydrolase. This enzyme also has base exchange activity towards NADP^+^ (Aarhus et al., 1995, Nam et al., 2020) to mediate the formation of NaADP, and can utilise drug-like cyclic substrates for the base exchange of NMN (Preugschat et al., 2008). We first investigated this using recombinant CD38 enzyme, to test whether it could mediate the conversion of amidated NAD^+^ precursors to their de-amidated counterparts, and vice versa. We used targeted mass spectrometry to measure the production of reaction products, including the putative base exchange product NaMN. One challenge to this approach is that NMN and NaMN are very close in molecular weight, and to overcome the issue of potential misidentification due to similar molecular weights, we utilised stable isotope labelled substrates containing ^13^C_6_ (M+6) or D_4_ tetra-deuterated (M+4) labelling of the nicotinyl ring (Fig. 2A), allowing greater mass separation for the clearer identification of newly formed base exchange products. If base exchange were to occur, incubation of NMN with isotope labelled nicotinic acid should result in the formation of labelled NaMN (Fig. 2A). To test this, unlabelled NMN was incubated with saturating levels of ^13^C_6_ nicotinic acid (M+6) in the presence of recombinant CD38 enzyme and/or the small molecule CD38 inhibitor **78c** (Haffner, Becherer et al., 2015) (Fig. 2B). If base exchange were to occur, this would result in the formation of M+6 labelled NaMN and unlabelled nicotinamide, though the formation of the latter product would also reflect the previously described NMN glycohydrolase activity of CD38 (Sauve et al., 1998). This reaction was tested under both acidic (pH 4) and neutral (pH 7) conditions, as the base exchange activity of CD38 towards NADP has been previously described to occur only within the acidified environment of endolysosomes (Aarhus et al., 1995, Nam et al., 2020). In line with our hypothesis, incubation with this enzyme reduced NMN levels and resulted in the formation of M+6 labelled NaMN (Fig. 2B) and unlabelled nicotinamide, although formation of the latter product also reflects its previously described NMN glycohydrolase activity. Unlike previously described base exchange activity for CD38 that only occurred under acidic conditions (Aarhus et al., 1995, Graeff, Franco et al., 1998), we detected NMN base exchange activity under both acidic (pH 4) and neutral (pH 7) conditions.

**Figure 2.**
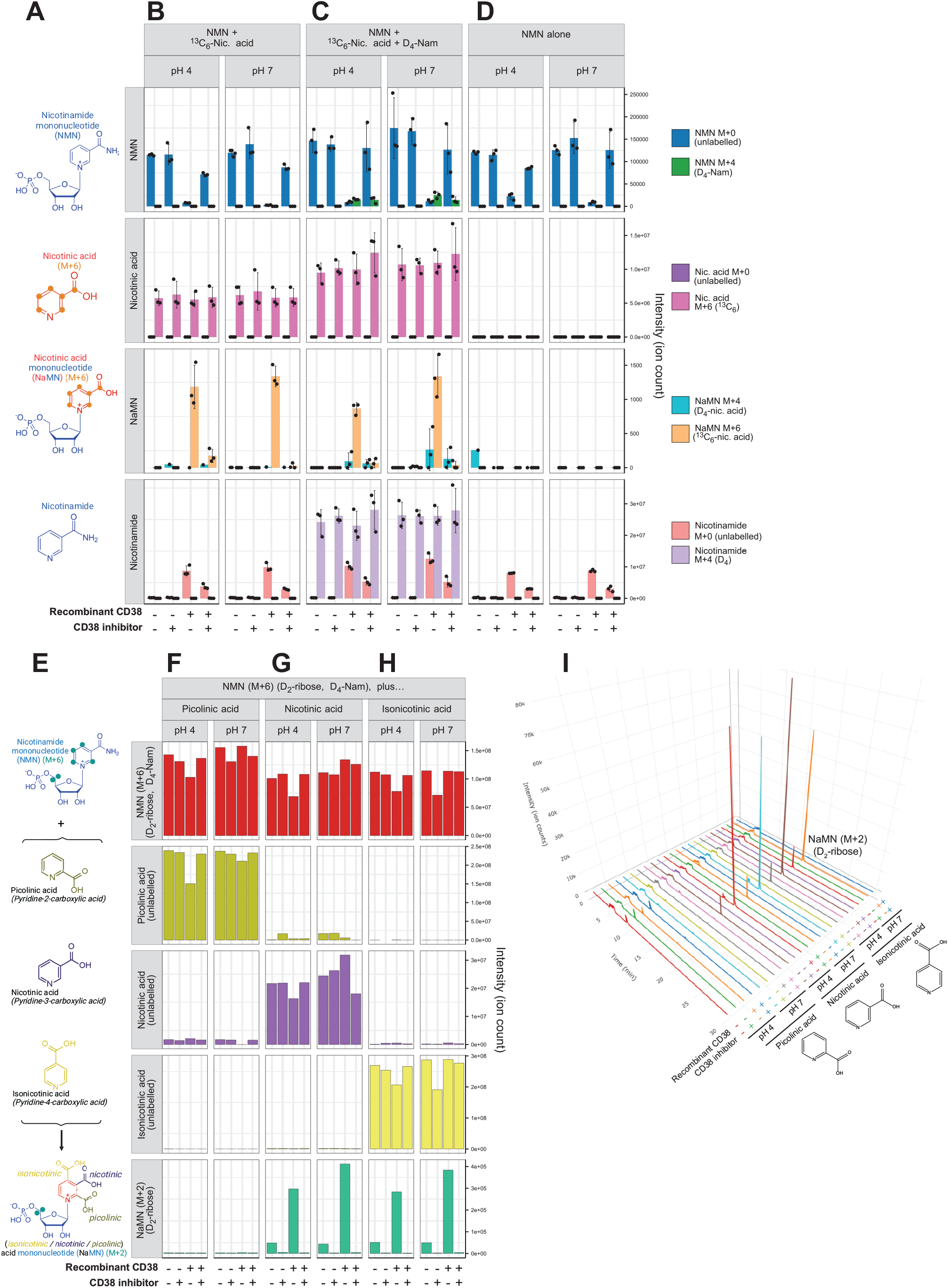
CD38 mediates NMN base exchange with nicotinic acid. A) Experimental scheme to identify NMN base exchange. Unlabelled NMN (M+0) and ^13^C_6_-labelled nicotinic acid (M+6) are co-incubated, and the formation of M+6 labelled nicotinic acid mononucleotide (NaMN) is indicative of base exchange of the unlabelled nicotinamide group from NMN for an M+6 labelled nicotinic acid. B) Targeted mass spectrometry for these metabolites in the presence or absence of recombinant CD38 protein, with or without co-treatment with the small molecule CD38 inhibitor **78c**, showing CD38-dependent formation of M+6 labelled NaMN. This base exchange is preferential for nicotinic acid over nicotinamide, as C) co-incubation of unlabelled NMN with both ^13^C_6_-nicotinic acid (M+6) and D_4_-nicotinamide (M+4) leads to the greater formation of M+6 NaMN over M+4 NMN. D) Technical control to show the absence of labelled NaMN in the absence of ^13^C_6_-nicotinic acid (M+6). E) Scheme to test whether CD38-mediated base exchange is selective to nicotinic acid. Double-labelled NMN containing D_2_-ribose and D_4_-nicotinamide, for an overall mass shift of M+6 was co-incubated with nicotinic acid or its structural orthologues picolinic acid or isonicotinic acid, which are identical in molecular weight. CD38 mediated base-exchange would result in exchange of the D_4_-nicotinamide label for an unlabelled group, resulting in the formation of a product with a molecular weight identical to NaMN plus an M+2 mass shift, due to retention of the ribose label. Base exchange was not observed for F) picolinic acid, but was observed for G) nicotinic acid and H) isonicotinic acid. I) Chromatograms showing the detection of M+2 ribose-labelled NaMN in each group. Error bars are standard deviations.

One possibility could be that CD38 mediates a non-selective base exchange, swapping in either a nicotinic acid or a nicotinamide, in the latter case as a form of “auto-base exchange” that would still yield NMN. To test whether this base exchange reaction was selective, this previous reaction (Fig. 2B) was repeated with the addition of D_4_-nicotinamide (M+4), which was present in equimolar amounts with ^13^C_6_ nicotinic acid (M+6), along with NMN (Fig. 2C). The formation of M+6 labelled NaMN was unaffected by the addition of D_4_-nicotinamide (M+4), however M+4 labelled NMN was also detected, albeit the formation of this product was not sensitive to the small molecule CD38 inhibitor **78c** (Fig. 2C). There was also a small amount of M+4 labelled NaMN, and while it is conceivable that CD38 could first mediate the auto-base exchange of NMN to yield M+4 labelled NMN, followed by some previously undescribed base exchange reaction, this product most likely reflects the detection of naturally occurring M+1 isotopes of NMN. These isotopes would have the same molecular weight as NaMN, and this emphasises the importance of our stable isotope labelling strategy in this investigation to better resolve the separation of these species. Finally, this reaction was also performed in the absence of base exchange substrates, with NMN alone (Fig. 2D). As expected, in this condition there was no detection of M+4 or M+6 labelled NaMN (Fig. 2D).

Next, we sought to further explore the selectivity of CD38 mediated base exchange between NMN and nicotinic acid by testing whether isomers of nicotinic acid would also act as base exchange substrates with NMN (Fig. 2E). To test this, we generated 5,5-D_2_-ribose, D_4_-nicotinamide labelled NMN, containing an M+2 label on the ribose group and an M+4 label on the nicotinamide group for an overall mass shift of M+6. This “double-labelled” NMN isotopologue should retain the M+2 ribose label following loss or exchange of the M+4 labelled nicotinamide group, which would result in a mass shift from M+6 to M+2. Double-labelled NMN (M+6) was co-incubated with unlabelled nicotinic acid, isonicotinic acid and picolinic acid (Fig. 2E), which contain the carboxylic acid side group at meta, para and ortho positions of the pyridine ring, respectively. Co-incubation of double-labelled NMN (M+6) with picolinic acid (pyridine-2-carboxylic acid) did not lead to the formation of M+2 ribose labelled NaMN (Fig. 2F), in contrast to incubation with unlabelled nicotinic acid (pyridine-3-carboxylic acid) (Fig. 2G), which was in line with the previous experiment (Fig. 2B-C). Interestingly, base exchange was also observed with isonicotinic acid (pyridine-4-carboxylic acid) (Fig. 2H). As in previous experiments (Fig. 2-C), the formation of M+2 labelled NaMN from both nicotinic acid and isonicotinic acid occurred at both pH 4 and pH 7 (Fig. 2G-H). For these latter two substrates, there was a small but detectable level of base exchange that occurred in the absence of CD38, indicating that this reaction can spontaneously occur in the absence of CD38. Interestingly, this spontaneous base exchange that occurred in the absence of CD38 was abolished by the presence of the small molecule inhibitor **78c** (Haffner et al., 2015). Representative chromatograms for the formation of this M+2 labelled NaMN product are shown in Fig. 2I, with the major product eluting at around 17 min (Fig. 2I).

A core insight that allowed for our discovery was the use of stable isotope labelling, as NMN and NaMN have similar spectral characteristics, similar elution times, and are almost identical in molecular weight, differing by only a single mass unit, with the signal from a lesser abundant NaMN potentially swamped by natural isotopes of NMN. One caveat to this approach is that heavy isotopes can impact enzyme function through kinetic isotope effects (KIEs). To ensure that this result was robust, and not an artefact of this particular isotope labelling combination, we next repeated this experiment using orthogonal labelling approaches, with three different combinations of isotope labels.

This included generating 5,5-D_2_-ribose NMN, with an M+2 label on the ribose group that should be retained following exchange of the nicotinamide group (Fig. 3A). This M+2 ribose labelled NMN was incubated with ^13^C_6_ nicotinic acid (M+6) (Fig. 3A). In the presence of recombinant CD38, there was a reduction in overall NMN levels, with production of M+8 labelled NaMN, indicating that M+2 labelled NMN had swapped its nicotinamide ring for a ^13^C_6_ nicotinic acid (M+6), liberating a free, unlabelled nicotinamide group (Fig. 3A). This single labelled NMN (M+2) preparation was found to be impure, containing a minor degree of unlabelled NMN (M+0), likely related to its enzymatic synthesis. As a result, there was also a lesser amount of M+6 labelled NaMN in samples that were co-incubated with CD38, reflecting base exchange of unlabelled NMN with ^13^C_6_ nicotinic acid (M+6) (Fig. 3A). Both groups contained M+2 labelled NaMN, however as described above, this likely reflects a signal from naturally occurring isotopes of NMN, as these two compounds differ by only single mass unit, with close elution times.

**Figure 3.**
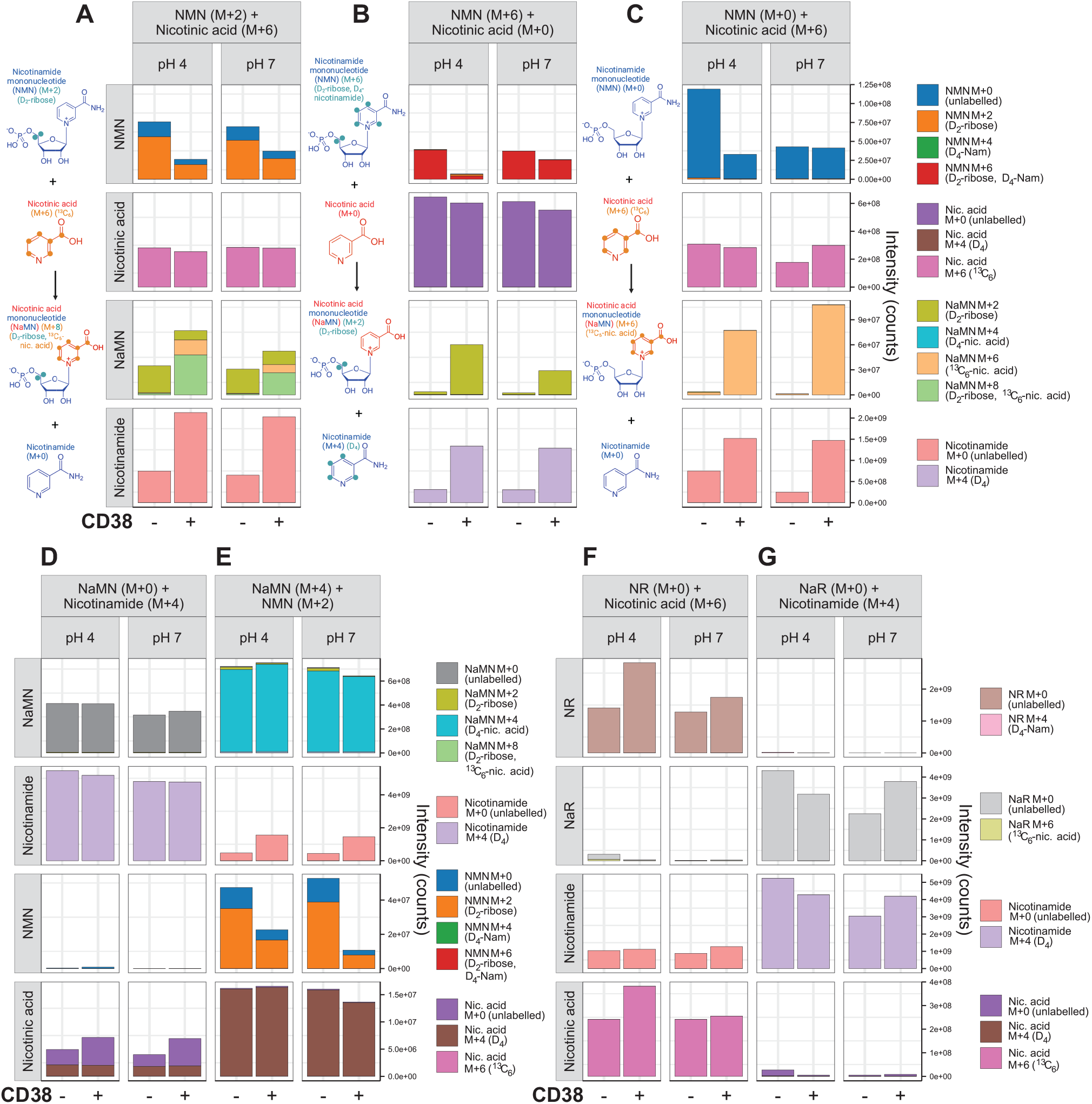
CD38 base exchange is selective to NMN, and not NaMN, NR or NaR. CD38 mediated base exchange was confirmed through three different labelling combinations. A) 5,5-D_2_-ribose labelled NMN (M+2) was co-incubated with ^13^C_6_-nicotinic acid, leading to the formation of D_2_-ribose, ^13^C_6_-nicotinic acid labelled NaMN (M+8). B) 5,5-D_2_-ribose, D_4_-nicotinamide labelled NMN (M+6) was co-incubated with unlabelled nicotinic acid, leading to the formation of D_2_-ribose labelled NaMN (M+2). C) Unlabelled NMN was co-incubated with ^13^C_6_-nicotinic acid, leading to the formation of ^13^C_6_-nicotinic acid labelled NaMN (M+6). D) To test whether CD38 could mediate base exchange on NaMN rather than NMN, unlabelled NaMN was co-incubated with D_4_-labelled nicotinamide (M+4), however M+4 labelled NMN was not identified. E) To test whether there is direct base exchange between NMN and NaMN, D_4_-nicotinic acid labelled NaMN (M+4) was co-incubated with D_2_-ribose labelled NMN (M+2), however the hypothetical base exchange products were not detected. F, G) To test whether nicotinamide riboside (NR) or nicotinic acid riboside (NaR) could act as a substrate for base exchange, F) unlabelled NR was co-incubated with ^13^C_6_-nicotinic acid (M+6), and G) unlabelled NaR was co-incubated with D_4_-nicotinamide (M+4), with no base exchange products observed in either case.

As in the previous experiment (Fig. 2E), we also used 5,5-D_2_-ribose, D_4_-nicotinamide labelled NMN, containing an M+2 label on the ribose group and an M+4 label on the nicotinamide group for an overall mass shift of M+6 (Fig. 3B). This double-labelled NMN (M+6) was incubated with unlabelled nicotinic acid in the presence or absence of recombinant CD38. This led to the presence of M+2 labelled NaMN, reflecting the loss of the M+4 labelled nicotinamide group from NMN, and its replacement with unlabelled nicotinic acid (Fig. 3B). Consistent with this, this reaction liberated free M+4 labelled nicotinamide, indicating its loss from M+6 double labelled NMN (Fig. 3B).

Finally, in line with our previous experiment (Fig. 2A), we again included the combination of unlabelled NMN with ^13^C_6_-nicotinic acid (M+6), again yielding M+6 labelled NaMN to indicate the exchange of the unlabelled nicotinamide ring of NMN with free ^13^C_6_-nicotinic acid (Fig. 3C). In each of these combinations, we consistently observed that base exchange activity was not impacted by pH, which is in contrast to the previously described base exchange activity of CD38 in the acidic endolysosomes for its production of NaADP from NADP (Aarhus et al., 1995, Fang et al., 2018, Graeff et al., 1998, Li & Wu, 2021). Together, these orthogonal approaches show that CD38 is capable of mediating base exchange on NMN with free nicotinic acid to yield NaMN, a deamidated intermediate of the Preiss-Handler / *de novo* pathways.

Having shown that CD38 could utilise nicotinic acid as a substrate for a base exchange reaction (Fig. 2B-C, 3A-C), we next tested whether this reaction could occur in the reverse direction, i.e. whether CD38 could carry out base exchange between NaMN and free nicotinamide to form NMN. Incubation of NaMN with D_4_-nicotinamide (M+4) did not result in the formation of M+4 labelled NMN (Fig. 3D), suggesting that this reaction only occurs in the direction of NMN to NaMN. While unlikely, we also tested whether the nicotinic acid of NaMN could be used as a substrate for NMN base exchange through using NMN that was labelled at its ribose group only (M+2), along with NaMN that was deuterated at the nicotinic acid group only (M+4) (Fig. 3E). If base exchange was to occur here, this should result in the formation of M+6 labelled NaMN. Unsurprisingly, this product was not observed (Fig. 3E). CD38 has strong glycohydrolase activity towards NMN (Sauve et al., 1998), however in each of these experiments when CD38 was incubated with NaMN, we did not observe any reduction in its levels, in contrast to reductions in NMN, suggesting that NaMN is resistant to the glycohydrolase activity of this enzyme (Fig. 3E). This observation could have important implications for the use of NAD+ precursors as therapeutics, whereby NaMN, unlike the more commonly used NMN, is more resistant to hydrolysis by CD38.

Finally, we also tested whether CD38 could mediate base exchange between nicotinamide riboside (NR) and nicotinic acid, as has previously been described for another cell surface enzyme, CD157/BST1 (Yaku et al., 2021). Unlike with NMN, when unlabelled NR was co-incubated with ^13^C_6_-nicotinic acid (M+6) in the presence of CD38, we did not observe M+6 labelled NaR (Fig. 3F). Similarly, when unlabelled nicotinic acid riboside (NaR) was co-incubated with D_4_-nicotinamide (M+4), we could not detect the production of M+4 labelled NR (Fig. 3G). Together, these data show that CD38 mediated base exchange between NMN and nicotinic acid to form NaMN is a new selective property of this enzyme, which does not carry out base exchange on NaMN, NR or NaR (Fig. 3D-G).

Next, we sought to test whether the demonstration of *in vitro* NMN base exchange activity (Fig. 2, 3A-C) was relevant to the formation of NaMN and NaAD following exogenous NMN treatment *in vivo*, as has been described in both humans and mice (Igarashi et al., 2022, Kim et al., 2023). We used NMN treatment in C57BL6 mice in a 2*2 experimental design, where animals received an acute oral bolus of NMN or the CD38 small molecule inhibitor **78c** (Haffner et al., 2015) or both compounds together (Fig. 4). Two hours later, animals were euthanased, tissues rapidly dissected and preserved for metabolite extraction and metabolomics analysis. In line with previous work (Igarashi et al., 2022, Kim et al., 2023), NMN treatment strongly increased levels of the deamidated metabolites NaMN and NaAD in liver and muscle, however this increase was abolished through co-administration of a CD38 inhibitor (Fig. 4). These findings (Fig. 4) helped to validate the physiological relevance of our *in vitro* discovery of the ability of CD38 to mediate the formation of NaMN from NMN through its base exchange activity (Fig. 2, 3A-C). Further, they provide a compelling mechanistic explanation for the previously described increase in the de-amidated metabolites NaMN and NaAD following exogenous administration with the amidated precursor NMN (Igarashi et al., 2022, Kim et al., 2023).

**Figure 4.**
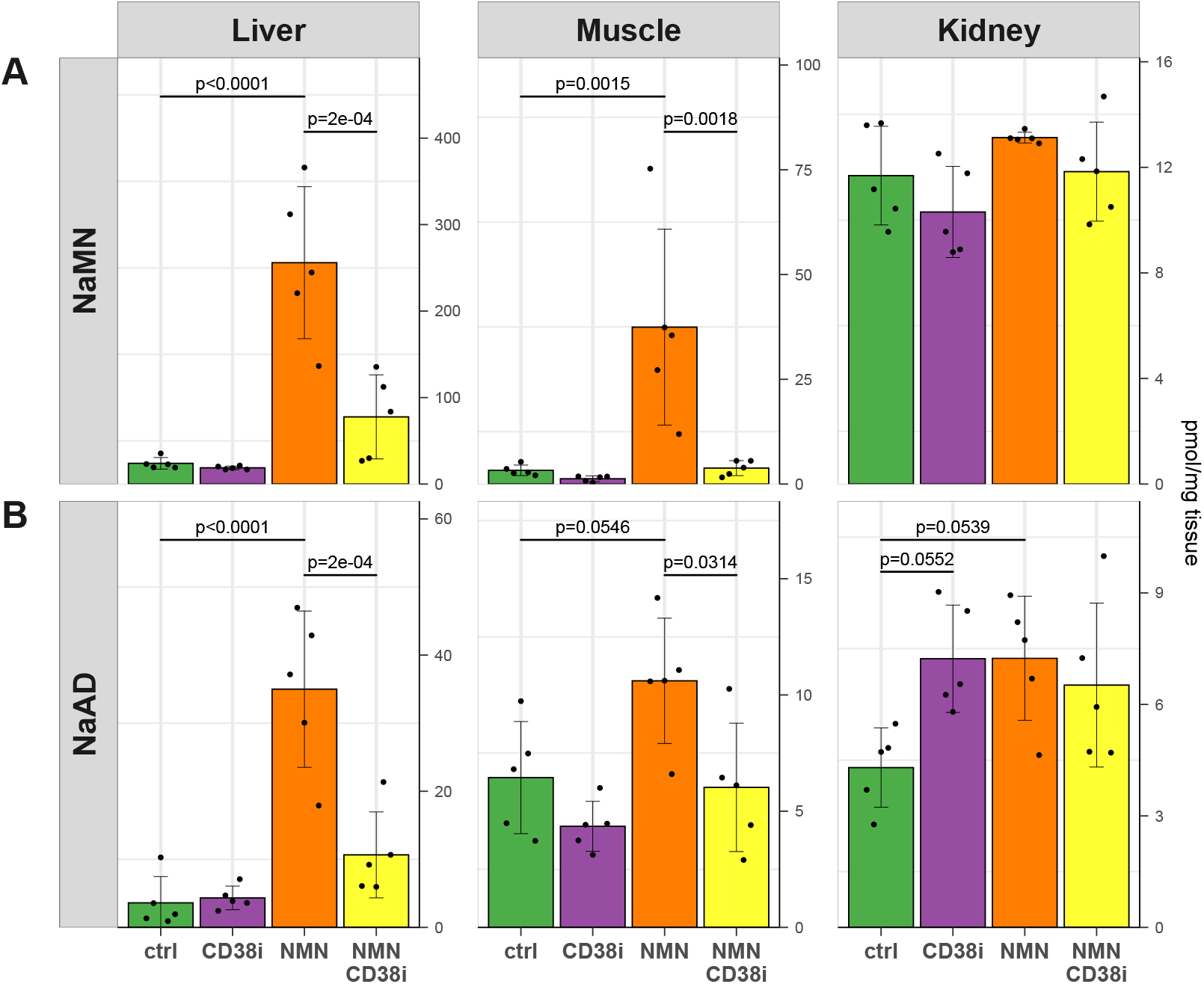
CD38 inhibition blocks NMN-induced increases in NaMN and NaAD in vivo. Mice received a single oral dose of the amidated NAD^+^ precursor NMN (500 mg/kg) in the presence or absence of the small molecule CD38 inhibitor 78c (“CD38i”) (Haffner et al., 2015) (10 mg/kg). Two hours later, animals were euthanased and liver, muscle (quadriceps) and kidneys were collected for metabolomics to measure levels of the Preiss-Handler / de novo pathway deamidated intermediates A) nicotinic acid mononucleotide (NaMN) and B) nicotinic acid adenine dinucleotide (NaAD). Data analysed by 2-way ANOVA followed by Tukey’s test for post-hoc comparisons, n=5/group, error bars are S.D.

## Discussion

The goal of this work was to identify additional possible pathways by which the amidated and deamidated arms of NAD^+^ biosynthesis could intersect in mammals. As with NMN, treatment with the amidated precursor NR also leads to a sharp increase in the deamidated NAD^+^ precursor NaAD (Trammell et al., 2016). Similar to our own work, this was recently shown to occur via a base exchange mechanism mediated by the cell surface enzyme CD157 / BST1, whereby the nicotinamide group of NR was exchanged for a free nicotinic acid to yield nicotinic acid riboside (NaR) (Yaku et al., 2021). This finding provided a strong mechanistic explanation for elevation of the deamidated arm of NAD biosynthesis following treatment with NR, however was exclusive to NR only, with no base exchange of CD157/BST1 observed towards NMN (Yaku et al., 2021) – an activity that we now show can be mediated by a different enzyme, CD38 (Fig. 1D).

While not investigated here, it would be interesting to know whether CD38 also contributes to the impact of NR on NaAD levels, as in addition to its base exchange by CD157/BST1 into NaR, NR is also converted to NMN as an intermediate *en route* to NAD^+^ biosynthesis (Bieganowski & Brenner, 2004). CD38 is a membrane bound protein, however its topology is not limited to facing the extracellular side of the plasma membrane: it can also protrude from the intracellular face of the plasma membrane, and is also present on intracellular vesicle membranes such as endolysosomes (Aksoy, White et al., 2006, Li & Wu, 2021, Zhao, Lam et al., 2012). It would be interesting to know whether exogenous NR can then undergo base exchange by intracellular CD38 following its conversion to NMN. Of relevance to this question, it would be interesting to know whether the physiological role of CD38 in mediating the impact of NMN treatment on NaMN and NaAD levels (Fig. 4) occurs due to extracellular or intracellular facing CD38. NMN enters the cell following its extracellular dephosphorylation into NR (Ratajczak, Joffraud et al., 2016), with some controversy around the existence of an intact NMN transporter (Grozio, Mills et al., 2019, Schmidt & Brenner, 2019, Wu & Sinclair, 2019). Assuming that NMN enters the cell in the form of NR and that CD38 mediated NMN base exchange is intracellular, it could be conceivable that CD38 might also play a role in the increase in NaMN and NaAD following NR treatment. This idea is however complicated by the evidence from Yaku et al, who show that the NR-induced spike in NaMN and NaAD is abolished in BST1 knockout mice (Yaku et al., 2021).

Another implication for the role of CD157/BST1 in mediating base exchange on NR (Yaku et al., 2021) is not only to explain the formation of the deamidated intermediate NaAD, but in explaining the results of isotope labelling studies. Isotope tracing studies performed by Trammell et al utilised double isotope labelled NR, containing stable isotope labels at both the ribose and nicotinamide groups (Trammell et al., 2016). When delivered *in vivo*, this double-labelled NR led to the formation of NaAD containing a single label only, along with predominantly single labelling of NAD^+^ (Trammell et al., 2016). There are several possible explanations for the observation of single, rather than double labelling. Firstly, once assimilated into NAD^+^, its rate of turnover (Liu et al., 2018) would lead to the separation of these labels, including the release of free nicotinamide that would be recycled back into NAD^+^ by the time tissues had been taken from animals following the initial NR bolus. Another possibility is that NR undergoes first pass metabolism in the gut and liver, being broken down into free nicotinamide prior to its cellular uptake and incorporation into the NAD^+^ metabolome (Chellappa et al., 2022, Liu et al., 2018). This alone might explain single labelling of NAD^+^, but not the formation of single labelled NaAD. Unlike mammals, bacteria in the gut microbiome express enzymes such as PncA, which can deamidate free nicotinamide into nicotinic acid, with the microbiome being responsible for the formation of nicotinic acid following administration with exogenous nicotinamide (Shats et al., 2020) and NR (Chellappa et al., 2022). If NR was being broken down into free nicotinamide, it is conceivable that its bacterial deamidation into nicotinic acid and assimilation into NAD^+^ through the Preiss-Handler pathway could account for the formation of single labelled NaAD. Indeed, it has been proposed that NR is assimilated following its recycling between the host and gut microbiome via nicotinic acid as an intermediate (Chellappa et al., 2022). The discovery that CD157/BST1 also acts as an NR base exchange enzyme (Yaku et al., 2021) could also explain the formation of single labelled, deamidated intermediates (i.e. NaAD) following double-labelled NR treatment (Trammell et al., 2016). It is, however, unclear as to what extent this base exchange mechanism is physiologically relevant, compared to first pass metabolism of NR into free nicotinamide, and its subsequent bacterial deamidation into nicotinic acid (Chellappa et al., 2022, Liu et al., 2018).

These questions related to NR metabolism are highly analogous to the findings presented here. As with NR, treatment with double labelled NMN *in vivo* leads to relatively low levels of double labelled NAD^+^ (Kim et al., 2023, Sauve, Wang et al., 2023). This could reflect the later timepoints used for those measurements, with intact NAD^+^ labelling likely to have undergone recycling (Liu et al., 2018) by that timepoint (Kim et al., 2023). It might also reflect its first pass metabolism in the gut and liver prior to uptake and incorporation (Chellappa et al., 2022, Liu et al., 2018). We had previously described that treatment with antibiotics to ablate the gut microbiome could shift the intact incorporation of double-labelled NMN into NAD^+^ to the salvage pathway, rather than the de-amidated, NADS-dependent Preiss-Handler / *de novo* pathway (Kim et al., 2023). While that work demonstrated a role for the gut microbiome in deamidating NMN prior to its intact incorporation into NAD^+^, the CD38-dependent base-exchange mechanism described here could also account for the observation of single labelling of NAD^+^, as the nicotinamide group of NMN could be swapped out for an unlabelled nicotinic acid prior to its incorporation into NAD^+^.

The degree to which base-exchange mechanisms contribute to the systemic metabolism of exogenous NR or NMN remains to be determined. While these data show a clear role for CD38 in the formation of the deamidated metabolites NaMN and NaAD following NMN treatment *in vivo* (Fig. 4), it is unclear to what degree flux through these intermediates contributes to overall NAD^+^ homeostasis. Future work should aim to address this, using time-resolved *in vivo* flux analysis of the NAD metabolism (Liu et al., 2018, McReynolds, Chellappa et al., 2021) in the context of CD38 inhibition and stable isotope labelled NMN, or in the context of CD157/BST1 deletion with labelled NR. Although adding another layer of difficulty to what would already be a demanding study, this work would be especially interesting if performed in the context of NADS knockout mice (Szot, Cuny et al., 2024), which could help to tease out the contributions of the deamidated pathway to the assimilation of NR and NMN into the NAD^+^ metabolome.

Another future experiment that would help to quantify the degree to which this base exchange mechanism is relevant to the assimilation of exogenous NMN into NAD^+^ could be to use NMN that is labelled at the ribose and/or phosphate groups only, and administer it in combination *in vivo* with labelled nicotinic acid in the presence or absence of a CD38 inhibitor. This could be an expensive experiment, due to the need for custom labelled NMN and a saturating quantity of stable labelled nicotinic acid, however if successful, would provide a quantitative answer as to what proportion of exogenous NMN is incorporated following this base exchange mechanism.

It may be that the CD38 mediated base exchange of NMN into NaMN accounts for only a small proportion of its activity. If so, this understanding still provides a mechanistic understanding for the sharp increase in the deamidated NAD^+^ precursors NaMN and NaAD, which could act as sensitive biomarkers (Trammell et al., 2016) for exogenous NAD^+^ precursor administration. Understanding whether this base exchange pathway plays a physiologically relevant role for overall NAD^+^ homeostasis could, however, answer a more important question for the field. The rationale for using NMN and NR is that they bypass the rate-limiting step in NAD^+^ biosynthesis, which is the conversion of nicotinamide to NMN by the enzyme NAMPT (Revollo, Grimm et al., 2004). If it is indeed the case that orally delivered NMN and NR undergo extensive first pass metabolism to yield free nicotinamide prior to its uptake and delivery to other tissues, one might expect that simply using free nicotinamide, as has long been used as nutrient fortification around the world, would be an equally effective strategy to raising NAD^+^ levels. Aside from having hypothetical bioequivalence, providing nicotinamide instead would also be much cheaper, given the costs of NR and NMN synthesis. Remarkably, these direct comparisons between different NAD^+^ precursors are largely lacking in animal models of disease (Li & Wu, 2021). While there has been an explosion in studies that test the efficacy of NMN and NR in disease models, they are rarely conducted against a group treated with nicotinamide. One study (Trammell et al., 2016) conducting a direct head-to-head comparison of NR against nicotinamide and nicotinic acid demonstrated differences in the pharmacokinetic profile of NAD^+^ formation, with NR treatment achieving a greater peak in NAD^+^ levels, along with differences in the formation of other NAD metabolites. One of the more convincing arguments that these precursors have superior efficacy as intact molecules is that NR treatment (Frederick, Loro et al., 2016, Mukherjee, Chellappa et al., 2017) and NMN treatment (Stromsdorfer, Yamaguchi et al., 2016, Wang, Zhang et al., 2017) can partially overcome the pathophysiological impacts of deletion of the rate-limiting enzyme NAMPT, which should not be possible if these precursors are degraded into free nicotinamide by first pass metabolism prior to their uptake (Chellappa et al., 2022, Liu et al., 2018).

Here, we propose that CD38-mediated base exchange of NMN into NaMN could reconcile the low levels of intact NMN incorporation into NAD^+^ with its apparent efficacy as an NAD^+^ precursor that can bypass the deletion of the rate limiting NAD biosynthetic enzyme NAMPT. Further, this mechanism could also explain the sharp increase in the deamidated metabolites NaMN and NaAD following NMN treatment. We propose that this is analogous to the base exchange mechanism of NR, which is instead mediated by CD157/BST1 (Yaku et al., 2021). Finally, this work expands our understanding of the role of CD38 in NAD^+^ homeostasis. This enzyme has received attention as a therapeutic target (Haffner et al., 2015) due to its role as an NAD^+^ glycohydrolase (Fig. 1B) that controls overall NAD^+^ levels (Camacho-Pereira et al., 2016, Chini et al., 2020, Tarrago et al., 2018), while also playing important signalling roles due to its base exchange activity, which yields the signalling intermediate NaADP (Aarhus et al., 1995, Chini et al., 1995,

Nam et al., 2020) (Fig. 1C). In this work, we show that this enzyme can also mediate base exchange on NMN to yield NaMN (Fig. 1D). Together, this provides a new mechanism by which the amidated and deamidated pathways of mammalian NAD homeostasis can intersect.

Limitations to this study include that we did not yet establish the precise enzyme kinetics of CD38-mediated base exchange on NMN. Establishing these will require detailed time-course experiments, including variations in substrate concentrations to test enzyme affinity. While this has been well established for other activities of CD38, for example its glycohydrolase activity towards NMN, these were able to utilise measurements based on changes in optical absorbance, allowing for higher throughput in sample number. In our case, we were not able to show differences in absorbance of fluorescence characteristics between each of the substrates and products of our reactions, forcing us to rely on targeted mass spectrometry with stable isotope labelled substrates. Resolution of each of these metabolites by mass spec also relied on chromatographic separation, which together required around 30 min in mass spec run time per individual sample, limiting our overall throughput due to cost and instrument time restraints. Regardless, future studies should aim to establish these parameters for the base exchange of NMN and nicotinic acid to NaMN, including Michaelis-Menten kinetics and K_cat_. It would be especially interesting to determine whether this newly discovered activity is sensitive to the abundance of other NAD^+^ metabolites, and whether the reaction can be limited by product inhibition. Finally, as described above, while we showed that CD38 played a key role in the formation of the deamidated metabolites NaMN and NaAD following *in vivo* administration with the amidated metabolite NMN, it would be interesting to determine the quantitative extent to which this base exchange activity contributed to the overall incorporation of NMN into NAD^+^.

## Methods

### Mass spectrometry

Metabolites were extracted from tissues using an extraction buffer containing 40:40:20 methanol, acetonitrile and 0.1N formic acid with the addition of 1 μM thymine-D_4_ as an internal standard. Briefly, snap-frozen tissue samples were crushed into a powder using a mortar and pestle with a small amount of liquid nitrogen sitting on a bed of dry ice. From this powder, 20 mg was weighed and 300 μL extraction buffer (previously cooled to -30C) was added. Samples were then lysed in a pre-cooled Precellys Cryolys at 6,500 rpm for 30 seconds, followed by a 20 second wait, and repeated twice more. Samples were then centrifuged for 10 min at 14,000 g at 4°C. Supernatants were collected and dried down in a vacuum centrifuge with no heat. Dried samples were reconstituted in 200 μL of a solution containing a 10:90 ratio of mobile phase A (20 mM ammonium acetate, pH 9) to mobile phase B (100% acetonitrile).

For data shown in Figs. 2 and 3, samples were run on a Thermo TSQ Vantage Triple-Stage Quadrupole mass spec, coupled to a Vanquish HPLC system (Thermo) using an InfinityLab Poroshell 120 HILIC-Z column (2.1 × 150 mm, 2.7 μm, Agilent catalogue number 673775-924). The injection volume was 5 μl, and the instrument was set to a dwell time of 30 ms, with a Q1 resolution of 0.7 and Q3 resolution of 1.2, with CID gas (mTorr) at 1.5, using SRM parameters shown in Table 1 below. The flow rate was 200 μl/min, with the percentage of solvent B set at 90% (0-2 min), 70% (13 min), 40% (16 -20 min), 90% (21-30 min). Metabolites of interest eluted from this ingredient as follows: NR at 17.2 min, NaMN at 17.09 min, NMN at 17.3 min, NaR at 12.96 min, nicotinamide at 2.24 min and nicotinic acid at 2.04 min.

**Table 1.** Selected reaction monitoring (SRM) parameters for the targeted detection of metabolites shown in Figs 2, 3.

| Compound | Start Time (min) | End Time (min) | Polarity | Precursor (m/z) | Product (m/z) | Collision Energy (V) | RF Lens (V) |
| --- | --- | --- | --- | --- | --- | --- | --- |
| NA_80_0_0 | 0 | 30 | Positive | 124.038 | 80.071 | 22.47 | 78 |
| NA_80_4_4 | 0 | 30 | Positive | 128.038 | 84.071 | 22.47 | 78 |
| NA_80_4_3 | 0 | 30 | Positive | 128.038 | 83.071 | 22.47 | 78 |
| NA_80_6_5 | 0 | 30 | Positive | 130.038 | 85.071 | 22.47 | 78 |
| NaR_124_0_0 | 0 | 30 | Positive | 256.087 | 124.042 | 10.91 | 41 |
| NaR_124_4_4 | 0 | 30 | Positive | 260.087 | 128.042 | 10.91 | 41 |
| NaR_124_6_6 | 0 | 30 | Positive | 262.087 | 130.042 | 10.91 | 41 |
| NaMN_124_0_0 | 0 | 30 | Positive | 336 | 123.929 | 14.28 | 53 |
| NaMN_124_2_0 | 0 | 30 | Positive | 338 | 123.929 | 14.28 | 53 |
| NaMN_124_2_2 | 0 | 30 | Positive | 338 | 125.929 | 14.28 | 53 |
| NaMN_124_4_4 | 0 | 30 | Positive | 340 | 127.929 | 14.28 | 53 |
| NaMN_124_4_2 | 0 | 30 | Positive | 340 | 125.929 | 14.28 | 53 |
| NaMN_124_6_4 | 0 | 30 | Positive | 342 | 127.929 | 14.28 | 53 |
| NaMN_124_6_6 | 0 | 30 | Positive | 342 | 129.929 | 14.28 | 53 |
| NaMN_124_8_6 | 0 | 30 | Positive | 344 | 129.929 | 14.28 | 53 |
| NAM_80_0_0 | 0 | 30 | Positive | 123.038 | 80.071 | 21.49 | 59 |
| NAM_80_4_4 | 0 | 30 | Positive | 127.038 | 84.071 | 21.49 | 59 |
| NAM_80_4_3 | 0 | 30 | Positive | 127.038 | 83.071 | 21.49 | 59 |
| NR_123_0_0 | 0 | 30 | Positive | 255 | 123 | 23.5 | 64 |
| NR_123_4_4 | 0 | 30 | Positive | 259 | 127 | 23.5 | 64 |
| NMN_123_0_0 | 0 | 30 | Positive | 334.962 | 123 | 14.89 | 138 |
| NMN_123_2_0 | 0 | 30 | Positive | 336.962 | 123 | 14.89 | 138 |
| NMN_123_4_4 | 0 | 30 | Positive | 338.962 | 127 | 14.89 | 138 |
| NMN_123_6_4 | 0 | 30 | Positive | 340.962 | 127 | 14.89 | 138 |

For data shown in Fig. 4, samples were run on a 1260 Infinity LC system using an Amide XBridge BEH column (100 × 2.1 mm, Waters Corp., USA) coupled to a QTRAP 500 mass spec (SCIEX), as described previously (Kim et al., 2023).

### Enzyme assays

CD38 base exchange activity was assessed *in vitro* using recombinant CD38 (Val54 – Ile300) with a C-terminal 6xHis tag produced in NS0-derived mouse myeloma cell line from R&D Systems (catalogue number RDS2404-AC), with activity of >2,500 pmol/min/μg, as measured by its ability to convert nicotinamide guanine dinucleotide (NGD^+^) to cyclic GDP-ribose. The CD38 small molecule inhibitor used here was 4-[[*trans*-4-(2-Methoxyethoxy)cyclohexyl]amino]-1-methyl-6-(5-thiazolyl)-2(1*H*)-quinolinone (CAS #1700637-55-3), as described as compound **78c** in Haffner et al (Haffner et al., 2015). Reactions were carried out in 30 mM Tris, which was adjusted to pH 4 or pH 7 as shown in each figure. NMN and NaMN isotopologues were obtained under custom synthesis from GeneHarbor Biotechnology (Hong Kong Science and Technology Park, Shatin, Hong Kong SAR, China), who utilise an enzymatic method for synthesis. We obtained ^13^C_6_-nicotinic acid (CLM-9954) and D_4_-nicotinamide (DLM-6883) from Cambridge Isotope Laboratories (Cambridge, MA, United States). Base exchange was assayed with a 1:5 ratio of base exchange substrate (e.g. NMN, NaMN, NR, NaR) to free base (e.g. nicotinic acid, nicotinamide). Reactions took place in 6 μl volumes, containing 250 mM sucrose, 10 μg/ml bovine serum albumin (BSA), 30 mM Tris buffer that was adjusted to either pH 4 or pH 7, with 0.2 mM base exchange substrate (e.g. NMN, NaMN, NR, NaR) and 1 mM free base (nicotinic acid or nicotinamide), with or without the CD38 inhibitor 78c at 20 μM, with or without recombinant CD38 (3 ng per 10 μl reaction). Reactions were allowed to proceed for 30 min at room temperature, and quenched through the addition of 50 uL of a 1:1 mix of mobile phase A (20 mM ammonium acetate, pH 9) and mobile phase B (acetonitrile). Reaction products were then dried down under vacuum centrifuge for 2 hr, prior to resuspension for mass spectrometry.

### Animal studies

All experiments were approved by the UNSW Animal Care and Ethics Committee (ACEC), which operates under the animal ethics guidelines from the National Health and Medical Research Council (NHMRC) of Australia. Animals had ad libitum access to standard chow diet (Gordon’s stock feeds, Yanderra, NSW Australia) and acidified drinking water, and were maintained on a 12 hr light/dark cycle at 22°C and 80% humidity, in individually ventilated cages. Male C57BL/6J mice were obtained from the Animal Resource Centre (ARC) in Perth, WA, and allowed to acclimatise for at least one week prior to experiments. At 9 weeks of age, animals received a single dose of NMN (500 mg/kg) and/or the CD38 inhibitor 78c (10 mg/kg), for a 2*2 study design. Both compounds were delivered by oral gavage, and 2 hr later animals were euthanased and tissues rapidly dissected and snap frozen for mass spectrometry.

To prepare the CD38 inhibitor **78c** for *in vivo* administration, we first prepared a solution of 4 mg/ml 78c in 20% Capsitol in milliQ water. This was adjusted to pH 3 using citric acid powder, heated to 80°C on a hot plate, with the addition of 5% Solutol (Kalliphor HS15), and stirred for 15 min. Following this, the solution was left in a sonicating water bath for 30 min, prior to its use on the same day. This was then delivered to mice by oral gavage at a dose of 10 mg/kg. NMN was prepared for oral gavage through freshly diluting dry NMN powder (GeneHarbor Biotechnology, Shatin, Hong Kong SAR, China) in phosphate buffered saline (PBS) prior to its oral gavage at a dose of 500 mg/kg.

### Statistics

Data (Fig. 4) were analysed by 2-way ANOVA followed by Tukey’s HSD post-hoc test for between treatment comparisons. Error bars in figures indicate standard deviations. Data were analysed in *R* and figures generated using the R packages *ggplot2, ggh4x* and *rstatix*. A detailed, annotated R script showing our analysis has been uploaded to our open data site, please see “data availability” section below.

## Data availability

Mass spectrometry RAW files, CSV files and annotated R scripts and data files allowing for reproducible data analysis and the recreation of figures presented here have been uploaded to the Mendeley Data server, reserved DOI 10.17632/k72zvgyst6.1 (<u>preview link</u>).

## Acknowledgments

We wish to dedicate this manuscript to the memory of Professor Anthony A. Sauve (1964 – 2022), an NAD^+^ biochemist at The Department of Pharmacology in the Weill Medical College of Cornell University, New York. We are grateful to have enjoyed his sense of humour and energy, and are especially thankful for his discussions of the work presented here. We also wish to acknowledge Sydney Analytical at The University of Sydney for the use of their mass spec facilities, in addition to their guidance and training. We would also like to acknowledge the Animal Services stae at UNSW for assistance with care of the animals used in this project.

## Author contributions

RM and JLB conducted experiments, analysed data. SEH conducted experiments, supervised experiments and edited the manuscript. LEQ optimised mass spectrometry methods, analysed and interpreted data, supervised experiments, and edited this manuscript. NT conceived of this project, designed and conducted experiments, obtained funding, supervised the project, edited this manuscript. LEW conceived of this project, designed experiments, obtained funding, supervised the project, analysed data, prepared figures, and wrote this manuscript.

